# Switching of metabolic programs in response to light availability is an essential function of the cyanobacterial circadian clock

**DOI:** 10.1101/090688

**Authors:** Anna M. Puszynska, Erin K. O’Shea

## Abstract

The transcription factor RpaA is the master regulator of circadian transcription in cyanobacteria, driving genome-wide oscillations in mRNA abundance. Deletion of *rpaA* has no effect on viability in constant light conditions, but renders cells inviable in cycling conditions when light and dark periods alternate. We investigated the mechanisms underlying this viability defect, and demonstrate that the *rpaA*^-^ strain cannot maintain appropriate energy status at night, does not accumulate carbon reserves during the day, and is defective in transcription of genes crucial for utilization of carbohydrate stores at night. Reconstruction of carbon utilization pathways combined with provision of an external carbon source restores energy charge and viability of the *rpaA*^-^ strain in light/dark cycling conditions. Our observations highlight how a circadian program controls and temporally coordinates essential pathways in carbon metabolism to maximize fitness of cells facing periodic energy limitations.

## Introduction

Organisms across kingdoms of life have evolved circadian clocks to temporally align biological activities with the diurnal changes in the environment. In the cyanobacterium *Synechococcus elongatus* PCC7942, the core circadian oscillator is comprised of the KaiA, KaiB and KaiC proteins (Nishiwaki et al., 2007; Rust et al., 2007), that generate oscillations in the phosphorylation state of KaiC. Time information encoded in the phosphorylation state of KaiC is transmitted to the transcription factor RpaA (Takai et al., 2006; Taniguchi et al., 2011; Markson et al., 2013) to generate circadian changes in gene expression (Markson et al., 2013). In continuous light, the expression of more than 60% of protein-coding genes in *S. elongatus* is regulated in a circadian manner (Vijayan et al., 2009) with two main phases of gene expression: genes peaking at subjective dawn or subjective dusk, where the term “subjective” refers to an internal estimate of time in the absence of external cues. Deletion of *rpaA* disrupts these rhythms in mRNA abundance and arrests cells in a subjective dawn-like transcriptional state, rendering them unable to switch to a subjective dusk-like expression program (Markson et al., 2013).

While it is clear that the circadian transcriptional program schedules timing of gene expression when the clock is free-running in the absence of changes in external light, it is unclear how circadian control of gene expression is used under more physiologically relevant conditions when light and dark periods alternate. Exposure to darkness restricts energy availability, creating unique metabolic demands in cyanobacteria which depend on sunlight for energy production through photosynthesis. Strikingly, the *rpaA*^-^ strain exhibits defects in cell growth and viability in cyclic, but not in constant light environments (Takai et al., 2006), suggesting an important role for circadian regulation of gene expression in alternating light/dark cycles.

We find that both accumulation and utilization of the carbon reserves required for energy production during periods of darkness are defective in the *rpaA*^-^ strain, and that correction of these defects restores viability. Our results provide insight into the role of the circadian program in enhancing fitness of cyanobacteria through coordination of central carbon metabolism with the metabolic demands imposed by periodic changes in the external environment.

## Results

As reported previously (Takai et al., 2006), in constant light wild type and *rpaA*^-^ cells grow at the same rate (Figure 1A, top panel); however, the *rpaA*^-^ strain is not viable when cultured in alternating light/dark conditions (Figure 1A, bottom panel and Figure 1 - figure supplement 1A). The *rpaA*^-^ strain rapidly loses viability when incubated in the dark (Figure 1 - figure supplement 1B), suggesting that the defect is induced by darkness and not by repeated light-to-dark and dark-to-light transitions. Complementation of the *rpaA*^-^ strain with *rpaA* expressed from an ectopic site in the genome fully restored viability (Figure 1 - figure supplement 1A and Figure 1 - figure supplement 1B). In wild type cells, expression of the *kaiBC* genes depends on RpaA and thus deletion of *rpaA* abrogates the function of the KaiABC oscillator (Takai et al., 2006). To establish whether the defect in viability during dark periods results from loss of Kai oscillator function or from loss of RpaA function, we performed viability experiments using the “clock rescue” strain background in which *kaiBC* expression is made independent of RpaA (Teng et al., 2013). We observed that the *rpaA*^-^ “clock rescue” strain phenocopies the *rpaA*^-^ strain and the isogenic *rpaA*^+^ strain phenocopies the wild type strain, demonstrating that the viability defect stems from the loss of RpaA function and not from loss of Kai oscillator function (Figure 1B and Figure 1 - figure supplement 1C). Since phosphorylation of RpaA is required for binding to target promoters (Markson et al., 2013), we tested whether a strain expressing a non-phosphorylatable mutant form of RpaA, RpaA D53A, is able to grow under light/dark cycles in the “clock rescue” background. We found that this strain is also inviable, indicating that the active DNA-binding form of RpaA is required for cell viability during exposure to light/dark conditions (Figure 1 - figure supplement 1D).

**Figure 1.**
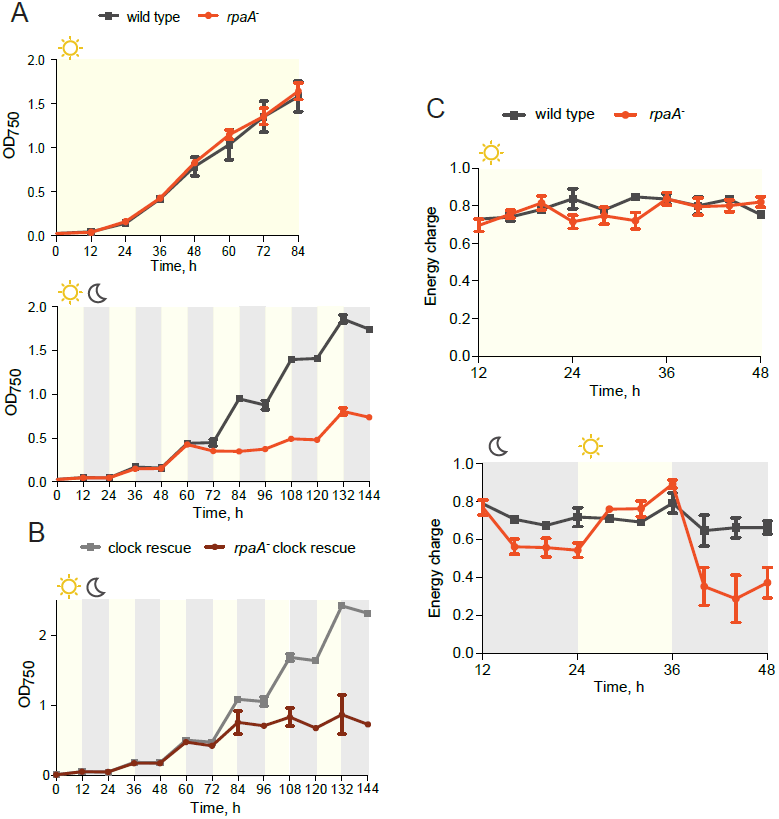
The *rpaA*^-^ strain displays a defect in viability and in maintenance of cellular energy levels in oscillating light-dark conditions. (A) Growth curves of wild type and the *rpaA*^-^ strain in constant light (top) and in 12 h light/12 h dark conditions (below) in BG-11 medium. (B) Growth curves of the “clock rescue” and the *rpaA*^-^ “clock rescue” strains in 12 h light/12 h dark conditions in BG-11 medium. Points represent the mean of three independent experiments with error bars displaying the standard error of the mean. (C) Changes in the energy charge, defined as ([ATP]+0.5*[ADP])/([ATP]+[ADP]+[AMP]), during growth of wild type and the *rpaA*^-^ strain in constant light (top) or in 12 h light/12 h dark conditions (bottom) in BG-11 medium. Each point represents the mean of three independent experiments with error bars displaying the standard error of the mean.

Circadian clocks and their output synchronize the metabolic activity of cells with diurnal fluctuations (Tu and McKnight, 2006; Green et al., 2008; Graf et al., 2010; Dodd et al., 2015). To understand if the observed viability defect stems from an inability of the *rpaA*^-^ strain (which is arrested in a ‘dawn-like’ transcriptional state (Markson et al., 2013)) to meet the metabolic requirements of darkness, we assessed cellular energy charge in the wild type and the *rpaA*^-^ strains over time in constant light conditions and in light/dark cycles. Energy charge describes the relative levels of ATP, ADP and AMP in the intracellular adenine nucleotide pool, and is a tightly controlled parameter that reflects the global metabolic energy status of the cell (Atkinson, 1968). In constant light, there was no difference in energy charge between the wild type and *rpaA*^-^ strains (Figure 1C, top panel). Wild type cells subjected to darkness experience only a slight reduction in energy charge (Figure 1C, bottom panel). In contrast, *rpaA*^-^ cells experience a reduction in energy charge during the first dark period and an even more dramatic reduction during the second dark period, suggesting that this strain has a defect in energy metabolism affecting energy maintenance during the dark phase of the light/dark cycle (Figure 1C, bottom panel). The decrease in energy charge was accompanied by a reduction in the total adenine nucleotide pool size in the *rpaA*^-^ strain (Figure 1 - figure supplement 2). A similar trend has been previously observed during starvation in carbon-limited cultures of *E. coli* (Chapman et al., 1971), prompting us to analyze changes in activity through pathways relevant for carbon metabolism in the strain lacking *rpaA*.

In constant light conditions, the circadian program directs expression of anabolic and catabolic pathways, with genes relevant for carbon catabolism peaking in mRNA abundance at subjective dusk (Vijayan et al., 2009; Diamond et al., 2015). To investigate the role of RpaA in orchestrating diurnal transcription of metabolic genes, we performed RNA sequencing in the *rpaA*^-^ “clock rescue” strain and an isogenic *rpaA*^+^ strain after exposure to darkness. The expression of circadian dusk-peaking genes was most affected in the *rpaA* deletion strain exposed to darkness (Figure 2 - figure supplement 1A). Specifically, we observed that genes encoding enzymes involved in glycogen breakdown, glycolysis and the oxidative pentose phosphate pathway (Figure 2 - figure supplement 1B and Supplementary File 1), which normally peak in expression at dusk, are among the genes with the greatest defect in expression in the *rpaA*^-^ cells (Figure 2A). These pathways are key for carbon utilization and energy production in the dark in cyanobacteria (Figure 2B), and their function is essential for survival of periods of darkness (Doolittle and Singer, 1974). Independent deletion of *gnd* as well as the *glgP_gap1* and *fbp_zwf_opcA* operons, which exhibit severely reduced expression in the *rpaA^-^* strain, results in impaired viability in light/dark cycles but not in constant light conditions (Figure 2 - figure supplement 1C and Doolittle and Singer, 1974; Scanlan et al., 1995). To confirm that the transcriptional defects in the *rpaA* deletion mutant affect carbon catabolism, we measured the enzymatic activities of glycogen phosphorylase (*glgP*), glucose-6-phosphate dehydrogenase (*zwf*) and 6-phosphogluconate dehydrogenase (*gnd*) in the dark, and observed that activity of each enzyme was strongly reduced in the *rpaA*^-^ strain compared to wild type (Figure 2C). Therefore, the *rpaA*^-^ strain appears to be unable to activate the carbon catabolic pathways upon the onset of darkness because it cannot induce transcription of the requisite enzymes at dusk.

**Figure 2.**
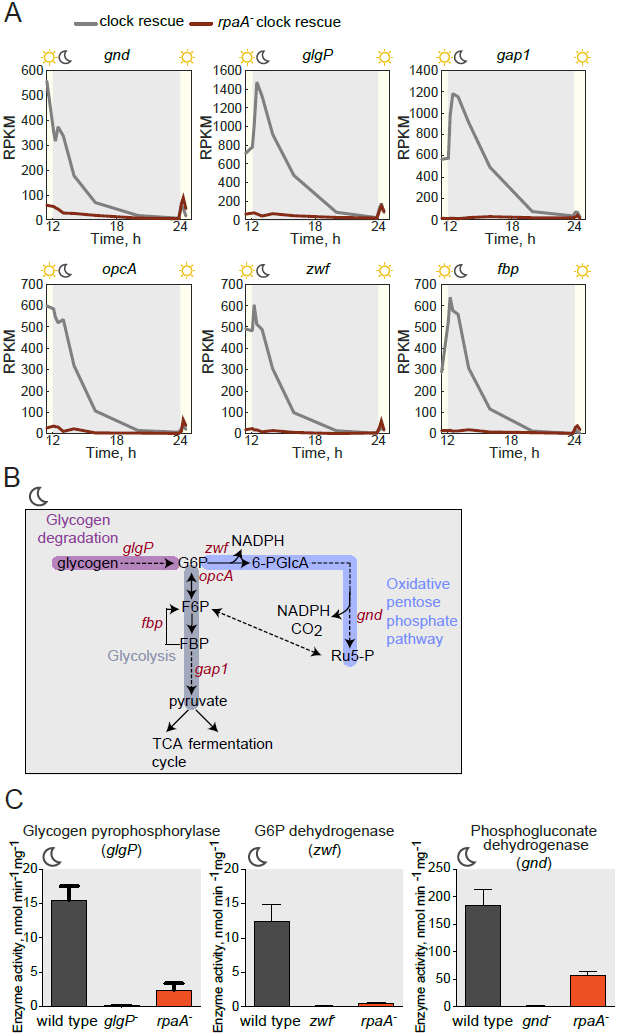
Sugar catabolism pathways are abrogated in the *rpaA^-^* strain. (A) Temporal expression profiles of genes encoding enzymes involved in sugar metabolism whose expression is abrogated in the *rpaA*^-^ “clock rescue” strain. Relative RNA levels were measured in the light/dark conditions by RNA sequencing. (B) A diagram representing carbon metabolism in *S. elongatus* PCC7942 in the dark. At night glycogen constitutes the store for energy and carbon skeletons. Glycogen degradation, glycolysis and oxidative pentose phosphate pathways are the key metabolic pathways operating at night. Deletion of operons with genes colored red leads to a strong viability defect specifically in light/dark conditions (Figure 2 - figure supplement 1C and Doolittle and Singer, 1974; Scanlan et al., 1995). *glgP*, glycogen phosphorylase; *gap1*, glyceraldehyde-3-phosphate dehydrogenase; *gnd*, 6-phosphogluconate dehydrogenase; *zwf*, glucose-6-phosphate dehydrogenase; *opcA*, glucose-6-phosphate dehydrogenase assembly protein; *fbp*, fructose-1,6-bisphosphatase, G6P, glucose-6-phosphate; 6-PGlcA, 6-phosphogluconate; F6P, fructose-6-phosphate; FBP, fructose-1,6-bisphosphate, Ru5-P, ribulose-5-phosphate (C) Enzymatic activities of glycogen phosphorylase, glucose-6-phosphate dehydrogenase and 6-phosphogluconate dehydrogenase in wild type, the *rpaA*^-^ and negative control strains measured 3 h after exposure to expected darkness. Error bars represent standard error of the mean of four independent experiments.

Glycogen is a crucial metabolic reserve used as a carbon and energy source during periods of darkness (Smith, 1983). In wild type cyanobacteria glycogen accumulates during the afternoon (Cervený and Nedbal, 2009; Knoop et al., 2010; Rugen et al., 2015) through the activity of a glycogen synthesis pathway composed of phosphoglucomutase (*pgm1* and *pgm2*), ADP-glucose pyrophosphorylase (*glgC*) and glycogen synthase (*glgA*) (Figure 3A). To analyze whether the dawn-arrested *rpaA*^-^ strain can accumulate glycogen reserves, we measured expression and activity of the glycogen synthesis enzymes and found that neither the expression levels nor activity of the enzymes in the pathway were significantly reduced in the *rpaA*^-^ strain (Figure 3B and 3C). Surprisingly, we observed that while glycogen content oscillates over a period of 24h in wild type cells both in constant light and in light/dark cycles, glycogen is present at a very low level in the *rpaA*^-^ strain regardless of light conditions, despite high levels of the synthetic enzymes in the extracts (Figure 3D). The same glycogen accumulation defect occurs in the *rpaA*^-^ “clock rescue” strain, demonstrating that it is a direct effect of deletion of *rpaA* (Figure 3E). Like the *rpaA*^-^ strain, glycogen synthesis mutants (*glgA*^-^ and *glgC*^-^) exhibit reduced viability in alternating light/dark conditions (Grundel et al., 2012, Figure 3 - figure supplement 1A, bottom panel) and in constant darkness (Figure 3 - figure supplement 1B), but grow somewhat slower than the *rpaA*^-^ strain in constant light conditions (Figure 3 - figure supplement 1A, top panel). We conclude that the deficiency in the preparation of a reserve carbon source – which occurs not at the level of transcription of the glycogen synthesis genes or regulation of the activity of the encoded enzymes – may contribute to the impaired viability of the *rpaA*^-^ strain in the dark.

**Figure 3.**
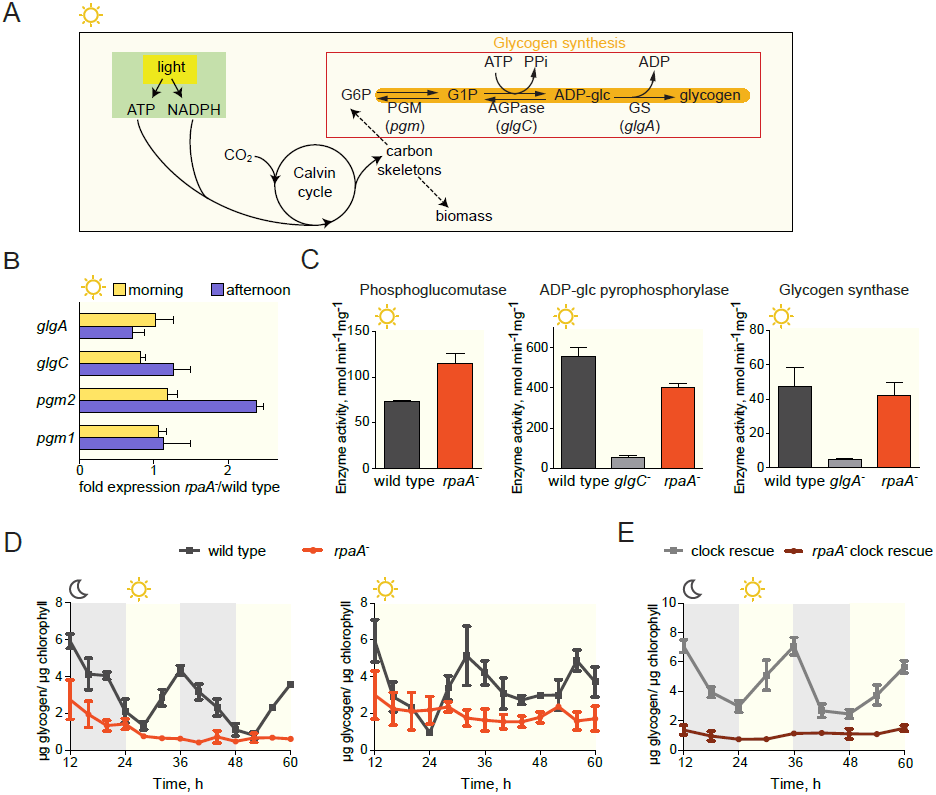
The *rpaA^-^* strain accumulates little glycogen. (A) A diagram representing carbon metabolism in *S. elongatus* PCC7942 in light. During the day *S. elongatus* cells perform photosynthesis to produce carbon skeletons and energy for growth and for preparation of glycogen stores. The glycogen synthesis pathway is comprised of three enzymatic activities: phosphoglucomutase (PGM), ADP-glucose pyrophosphorylase (AGPase) and glycogen synthase (GS). G6P, glucose-6-phosphate; G1P, glucose-1-phosphate; *pgm*, phosphoglucomutase; *glgC*, ADP-glucose pyrophosphorylase; *glgA*, glycogen synthase. (B) Relative expression of genes encoding enzymes in the glycogen synthesis pathway measured by RT-QPCR in the morning and in the afternoon in the wild type and *rpaA*^-^ strains. Error bars represent the standard error of the mean of three independent experiments. (C) Activities of enzymes in the glycogen synthesis pathway in wild type, the *rpaA*^-^ and negative control strains measured during the day (at time =10h). Error bars represent the standard error of the mean of three independent experiments. (D) Glycogen content in wild type and *rpaA*^-^ strains grown in light/dark and constant light conditions. Points represent the mean of two experiments with error bars displaying the standard error of the mean. (E) Glycogen content in the “clock rescue” and the *rpaA*^-^ “clock rescue” strains grown in light/dark conditions. Points represent the mean of two experiments with error bars displaying the standard error of the mean.

We hypothesized that the inability of the *rpaA*^-^ strain to maintain appropriate energy levels and cell viability in the dark stems from the defects in accumulation of carbon/energy stores during the day and their utilization at night. To assess whether restoration of the correct carbon catabolic route in darkness is sufficient to rescue the energy charge and viability of the *rpaA*^-^ strain in light/dark conditions, we reconstituted the activities of the carbon utilization pathways and provided a transporter for uptake of external carbon. We restored expression of the carbon utilization enzymes in the *rpaA*^-^ strain by placing their expression under an IPTG-inducible ectopic P*trc* promoter, and confirmed that induction with IPTG restored enzyme activity in cell extracts (Figure 4 - figure supplement 1). To counterbalance the inability of the *rpaA*^-^ cells to accumulate internal carbon reserves, we expressed the GalP transporter to allow glucose uptake from the medium. Expression of either the missing enzymes or the GalP transporter alone with supplementation of glucose in the *rpaA*^-^ background was not sufficient to restore viability of this strain (Figure 4A and 4B). However, restoration of the carbon utilization pathways combined with the introduction of the glucose transporter in the presence of glucose rescued the viability defect of the *rpaA*^-^ strain (Figure 4A and 4B). In this engineered strain, glucose feeding does not increase accumulation of internal carbon stores (Figure 4C), but does restore high energy charge levels in the *rpaA*^-^ cells (Figure 4D). Our results strongly suggest that functional circadian output mediated by the activity of RpaA is required for the anticipatory accumulation of carbon reserves during the day, as well as metabolic switching to carbon catabolism at dusk. These metabolic adaptations allow cells to stimulate energy production and sustain cell viability during periods of darkness that challenge and constrain resources for cyanobacterial growth.

**Figure 4.**
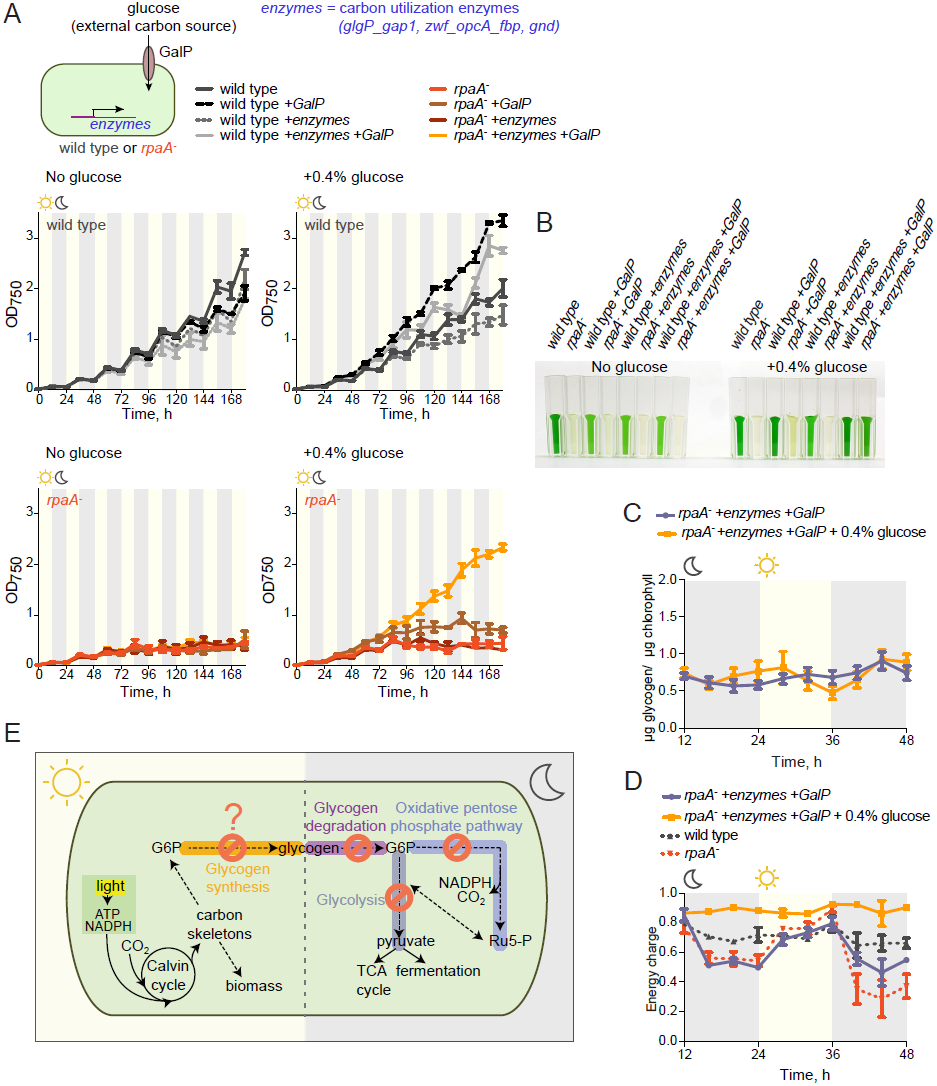
Restoration of sugar catabolism pathways and glucose feeding rescue the viability defect of the *rpaA*^-^ strain. (A) Growth curves of the indicated engineered strains in 12 h light/12 h dark conditions in BG-11 medium with 100 μM IPTG and with (right) or without (left) 0.4% glucose supplementation. Wild type (top) and the *rpaA*^-^ (below) strains were engineered to express a sugar transporter GalP and/or sugar utilization enzymes using an IPTG-inducible promoter. Points represent the mean of two experiments with error bars displaying the standard error of the mean. (B) Representative cell cultures of the engineered strains photographed at the end of the growth experiment in light/dark conditions with or without 0.4% glucose supplementation. *S. elongatus* cells are blue-green in color. The darker the color of the culture, the higher is its optical density at 750 nm. (C) Glycogen content in in the *rpaA*^-^ + *galP* + *enzymes* strain +/- 0.4% glucose in light/dark conditions. Points represent the mean of two experiments with error bars displaying the standard error of the mean. (D) Energy charge measurement in the *rpaA*^-^ + *galP* + *enzymes* strain with or without 0.4% glucose supplementation in light/dark conditions. Points represent the mean of two experiments with error bars displaying the standard error of the mean. Data representing the energy charge in wild type and the *rpaA*^-^ strain are reproduced from Figure 1C to facilitate comparison. (E) A model representing physiological processes in *S. elongatus* regulated by the activity of the circadian system that contribute to cell fitness. The circadian clock schedules periods of activity of RpaA to coordinate anticipatory carbon reserve formation during the day with carbon utilizing metabolic pathways activated at dusk to fuel cell integrity and viability in an environment in which light and dark periods alternate. Defects present in the *rpaA* deficient cells, indicated by red symbols on the diagram, lead to a low energy charge and reduced viability during periods of darkness.

## Discussion

Daily environmental changes in light availability give rise to periodic demand for metabolism of alternate energy sources in photosynthetic organisms (Doolittle, 1979). To maximize fitness, photosynthetic organisms need to accurately allocate resources by carrying out carbon assimilation during the day and utilizing stored carbohydrate reserves at night (Smith, 1983). Here we show that in the cyanobacterium *S. elongatus* PCC 7942, the transcription factor, RpaA, which orchestrates global circadian output, is critical for the coordination of carbon anabolism and catabolism and thus cell viability in light/dark cycles. Deletion of *rpaA* prevents glycogen accumulation in light and renders cells unable to utilize existing glycogen at dusk (Figure 4E) – this leads to alterations in cellular energy charge and, as previously observed, to a severe reduction in fitness. It is yet to be determined whether RpaA is rewiring carbon metabolism solely through transcriptional control or through other mechanisms.

The mechanism underlying perturbed carbon accumulation in the *rpaA*^-^ strain is unclear. The activities of glycogen synthesizing enzymes in the *rpaA*^-^ strain are high *in vitro*, however they could be affected *in vivo* by the redox state of the cell or by metabolite levels. It is also possible that an RpaA-dependent transcript expressed in the afternoon is required to funnel assimilated carbon into glycogen synthesis. Alternatively, additional metabolic flux changes may occur in the *rpaA*^-^ strain that prevent formation of glycogen stores.

Our data support a model in which anticipation of periodic dark-induced resource limitation is one essential role of the cyanobacterial circadian system. The output of the core oscillator, through regulation of RpaA activity, temporally coordinates metabolism to prepare cells for the metabolic demands of periods of darkness. Correct scheduling of central carbon metabolic activities allows cells to survive the night. Recently, Lambert et al. observed that the circadian clock orchestrates a trade-off between rapid cell growth in the morning and robustness to starvation in the afternoon, suggesting preparation for the night as a plausible role for the circadian clock (Lambert et al. 2016). Our findings are consistent with this model and provide novel mechanistic insight into circadian regulation of cyanobacterial energy metabolism. It remains to be investigated whether the circadian-driven preparation for darkness is a shared strategy employed by other photosynthetic organisms.

Further, our results suggest a molecular explanation for the observed selective advantage of circadian resonance. When the periods of the endogenous circadian system and periods of the environmental fluctuations are equal, organisms show a fitness advantage and outcompete mutants whose clocks have an altered periodicity in the same environment (Ouyang et al., 1998; Woelfle et al., 2004). The mechanistic basis for this observation has been lacking. Based on our findings, we expect that the fitness defects in the strains with core oscillator periods unequal to the period of environmental variation result from mistiming of RpaA activation. The resultant mistiming of accumulation and mobilization of glycogen stores with the onset of the nightfall would lead to a carbon deficit at night, and impact growth as in the *rpaA*^-^ mutant during dark periods. One role of the KaiABC clock in our model, therefore, is to correctly phase the activity of RpaA and downstream carbon metabolism with the external environment, maximizing fitness and growth.

Finally, our data together with a recent study by Diamond and colleagues (Diamond et al., 2015) emphasize the role of the circadian system in driving oscillations in metabolism. On the other hand, it has been demonstrated that metabolic processes provide feedback to the circadian clock by regulating its resetting, indicating an important role of the fluctuating metabolic status of a cell in driving circadian oscillations (Rust et al., 2011; Pattanayak et al., 2014; Pattanayak et al., 2015). Together, a complex bidirectional relationship between the circadian program and metabolism emerges in which the circadian system both controls and is controlled by the metabolic rhythms to set and tune cell physiology with the environmental cycles. This tight coupling between circadian machinery and metabolism is coming to light as a universal feature across kingdoms of life (Green et al., 2008; Dodd et al., 2015).

## Experimental Procedures

### Cyanobacterial strains

Strains were constructed using standard protocols for genomic integration by homologous recombination (Clerico et al., 2007) and are listed in Supplementary File 2. All plasmids were constructed using a protocol for Gibson assembly (Gibson et al., 2009).

The *rpaA*^-^ strain was constructed by transforming wild type cells with pΔRpaA(Gmr) that was a gift from Joe Markson. The “clock rescue” strain was constructed following Teng et al., 2013 by replacing the native P*kaiBC* promoter between base pairs 1240468 and 1240554 on *S. elongatus* chromosome with a P0050 promoter encompassing 508 base pairs upstream of the translation start of *synpcc7942_0050*. We used a kanamycin resistance cassette upstream of the promoter as a selection marker. The *rpaA*^-^ “clock rescue” strain was made by transforming pΔRpaA(Gmr) plasmid into the “clock rescue” strain background. The *rpaA*^-^ +*rpaA* strain was made by transforming wild type *S. elongatus* simultaneously with pΔRpaA(Gmr) and a NS 1 targeting vector pAM1303 carrying the *rpaA* gene with its native promoter encompassing 400 base pairs upstream of the translation start. The pAM1303 plasmid was a gift from Susan Golden. The *rpaA^-^ + empty plasmid* was created by transforming wild type cells with a pΔRpaA(Gmr) plasmid and an empty NS 1 targeting vector pAM1303. The “clock rescue” *_c_rpaA* and “clock rescue” *_c_rpaA D53A* strains were both made using the “clock rescue” strain background. In the *_c_rpaA* strains the *rpaA* promoter was changed to a P*0050* promoter allowing for continuous expression of *rpaA* and *rpaA D53A*. A gentamycin resistance cassette was placed upstream of the P*0050* promoter for selection. The D53A mutation was introduced following Markson et al., 2013.

The strain lacking the *glgP_gap1* (Synpcc7942_0244-0245) operon was made using the pBR322 plasmid carrying the kanamycin resistance cassette flanked by 1000 nucleotides of DNA from upstream and downstream of *glgP_gap1* locus. In the *zwffbp^-^opcA^-^* strain the *zwf_fbp_opcA* operon (Synpcc7942_2333-2335) was replaced with a kanamycin resistance cassette. In the *gnd^-^* strain the *gnd* gene (Synpcc7942_0039) was replaced with a kanamycin resistance cassette. The strain lacking *glgA* (Synpcc7942_2518) and the strain lacking *glgC* (Synpcc7942_0603) were made by using the pBR322 plasmids carrying the kanamycin resistance cassette flanked by 1000 nucleotides of DNA from upstream and downstream of respectively *glgA* and *glgC* locus.

The *wild type +galP* strain was made by transforming wild type cells with P*trc::galP* construct in a modified NS 1 targeting vector pAM1303 in which the cassette providing resistance to spectinomycin and streptomycin was replaced with a nourseothricin resistance cassette (Taton et al., 2014). The gene *galP* was amplified from the *E. coli* genomic DNA. The *wild type +enzymes* strain was made by sequentially replacing P*gnd*, P*glg_gap1* and P*zwf_fbp_opcA* with the P*trc* promoter making expression of *gnd, glgP_gap1* and *zwf_fbp_opcA* transcripts IPTG-inducible. The *wild type +enzymes +galP* strain was made by transforming the *wild type +enzymes* strain with P*trc::galP* construct in a modified NS 1 targeting vector pAM1303 that carried a nourseothricin resistance cassette. The *rpaA^-^ +galP, rpaA^-^ +enzymes* and *rpaA^-^ +enzymes +galP* strains were made by transforming pDRpaA(Gmr) plasmid respectively into *wild type +galP, wild type +enzymes* and *wild type +enzymes +galP* strains.

## Cell Culture

Cell cultures of wild type and mutant cells were grown under illumination with cool fluorescent light at 40 μE m^-2^ s^-1^ (μmoles photons m^-2^ s^-1^) in BG11 medium at 30 °C. For the light/dark experiments, cultures were incubated under alternating 12 h light/ 12 h dark conditions with the same light intensity of 40 μE m^-2^ s^-1^ during light periods. For experiments performed in Figure 4 strains were grown in BG-11 medium with 100 μM IPTG with or without supplementation with 0.4 % (w/v) glucose.

For the experiments described in Figure 1C and Figure 1 - supplement 2, Figure 2, Figure 3, and Figure 4 - figure supplement 1, wild type and “clock rescue” strains were entrained by exposure to 12 hours of darkness, followed by 12 h of light, followed by another 12 h of darkness. The *rpaA*^-^ and *rpaA*^-^ “clock rescue” strains were entrained by one pulse of 12 hours of darkness, followed by incubation in light to allow cell growth to resume. For the experiments described in Figures 4C and 4D the strains were entrained by one pulse of 12 hours of darkness and then incubated in light until the start of the experiment.

### Growth and viability assays

For the liquid growth assays, liquid cultures of wild type and mutant cells were pre-grown in continuous light in a medium lacking antibiotics. Cultures were diluted to OD_750_ = 0.02 and grown either in constant light or in 12 h light/12 h dark cycling conditions at 30 °C. Optical density of cells was monitored at OD_750_. Experiments were performed in duplicate (Figure 3 - figure supplement 1A, Figure 4A and 4B) or in triplicate (Figure 1A and 1B).

For the spot plate growth assays, liquid cultures of wild type and mutant cells were pre-grown in continuous light in a medium lacking antibiotics. Cells were diluted to OD_750_ = 0.25 and a dilution series was performed from 10^0^ to 10^-4^. Then, 10 μl of each dilution step were spotted onto BG11 agar plates with no antibiotics. Plates were incubated in constant light at 30 °C for 7 days or under 12 h light/12 h dark alternating conditions for 14 days. Experiments were performed in triplicate.

Viability after prolonged dark treatment was assessed by a colony forming unit assay. Light-grown cultures were diluted to OD_750_ = 0.025, transferred to darkness and sampled every 24h. Each aliquot removed from the culture was diluted in BG11 medium 1000 times by serial dilution. 100 μl of the diluted culture were plated onto a BG11 plate. Plates were incubated for 7 days at 30°C under constant illumination and colonies on each plate were counted. Viability was expressed as: % viability = N/N_0_ x 100%, where N_0_ is the colony count before exposure of cultures to darkness. Each experiment was performed in duplicate.

### Adenine nucleotide analysis

Nucleotides were extracted following the method used by Rust et al., 2011 with modifications. 2 ml of the cyanobacterial culture was added to 0.5 ml of ice-cold 3 M perchloric acid with 77 mM EDTA, vortexed briefly and incubated on ice for 5 min. The mixture was neutralized with 1.34 ml of 1 M KOH, 0.5 M Tris, 0.5 M KCl and then centrifuged at 4000 rpm for 20 min at 4 °C. The supernatant was then filtered through Amicon Ultra-4 filters (Millipore) and stored at –80 °C.

To measure adenine nucleotides, extracts were thawed and diluted 2.6X. For ATP measurement, extracts were diluted in a buffer containing 25 mM KCl, 50 mM MgSO_4_ and 100 mM HEPES, pH 7.4 (L buffer) with 1 mM phosphoenolpyruvate. To measure ADP + ATP, extracts were diluted in the above buffer containing also 3 U ml^-1^ type II pyruvate kinase from rabbit muscle (Sigma-Aldrich). To measure AMP + ADP + ATP, extracts were diluted in the L buffer containing 1 mM phosphoenolpyruvate, 3 U ml^-1^ type II pyruvate kinase from rabbit muscle and 75 U ml^-1^ myokinase from rabbit muscle (Sigma-Aldrich). Diluted extracts were incubated for 30 minutes in a 37 °C water bath followed by 10 minutes of heat treatment at 90 °C to inactivate enzymes. Extracts were assayed in triplicate in a black 96-well plate. 30 μl of the L buffer containing 35 μg ml^-1^ firefly luciferase (Sigma-Aldrich) and 1 mM luciferin (Sigma-Aldrich) were added to 260 μl of the extract. The luminescence signal was measured from each well in a TopCount luminescence counter (Perkin-Elmer).

## RNA-Seq

For the RNA-sequencing experiment the “clock rescue” and the *rpaA*^-^ “clock rescue” strains were entrained and harvested by filtering and freezing in liquid-nitrogen at following time points: dusk (ZT=12h); 5 min, 15 min, 30 min, 1 h, 2 h, 4 h, 8 h and 11 h 50 min after exposure to darkness; and 5 min, 15 min and 30 min after re-exposure to light at dawn.

RNA purification and library preparation were performed as described in Markson et al. 2013. NCBI reference sequences NC_007604.1, NC_004073.2, and NC_004990.1 were used to align sequencing reads to the *S. elongatus* chromosome and the endogenous plasmids with Bowtie. Uniquely mappable reads with maximum of three mismatches per read were allowed to map to the genome.

To quantify gene expression we counted the number of coding reads between the start and stop positions of open reading frames. We performed RPKM normalization and searched for differentially expressed genes that are at least 3-fold lower in the *rpaA*^-^ “clock rescue” strain than in the control in at least 5 of the 12 measured time points. We performed an analysis of the functional annotations of the protein coding genes whose expression is defective in the *rpaA*^-^ “clock rescue” strain using gene functions available in the Cyanobase. The sequencing data reported in this article can be found under the accession number GEO: xxxx.

## RT-qPCR for Gene Expression

Following entrainment (two cycles of entrainment for wild type strain and one cycle for the *rpaA*^-^ strain) equal ODs of wild type and the *rpaA*^-^ cells were harvested by filtering and freezing in liquid nitrogen in the morning (ZT=3h) and in the afternoon (ZT=9h). RNA extraction was performed as above. RT-qPCR was carried out using SYBRGreen PCR Master Mix (Applied Biosystems), Superscipt III Reverse Transcriptase (Invtrogen), and the following primers:

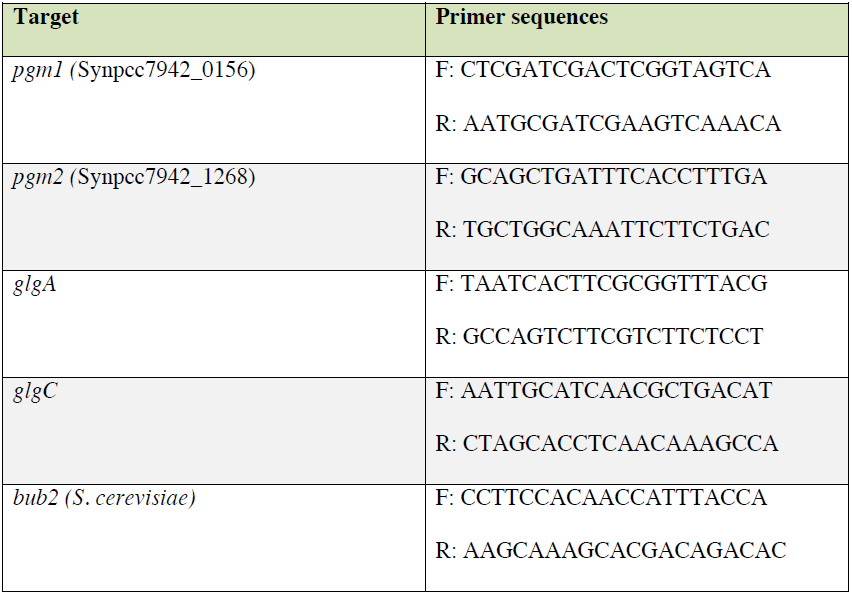

The abundance of transcripts was normalized by an external spike-in *bub2* transcript from *S. cerevisiae. bub2* was in vitro transcribed, added to the RNA AE extraction buffer and extracted together with *S. elongatus* RNA.

### Enzyme activity assays

All enzyme assays were performed using fresh cultures harvested immediately before the assay. Cells were collected by centrifugation. To prepare lysates cells were resuspended in an appropriate assay buffer as described below with Complete EDTA-Free Protease Inhibitors (Roche), transferred to 2 ml screw-cap tunes containing 0.1 mm glad beads (Research Products International Corp) and lysed by ten 30 sec long cycles of bead-beating at 4 °C with periodic cooling on ice. Protein concentration of each sample was determined by Bradford assay (Bio-Rad). Assays were performed in 96-well plate in the reaction volume of 200 μl. Glycogen phosphorylase activity was assayed in a buffer containing 18 mM KH_2_PO_4_, 27 mM Na_2_HPO_4_, 15 mM MgCl_2_ and 100 μM EDTA with 340 μM Na_2_NADP^+^, 4 μM glucose-1,6-bisphosphate (Sigma-Aldrich), 0.8 U ml^-1^ phosphoglucomutase (Sigma-Aldrich), 6 U ml^-1^ glucose-6-phosphate dehydrogenase (Sigma-Aldrich) and 2 mg ml^-1^ glycogen from bovine liver (Sigma-Aldrich) as substrate (Fu and Xu, 2006). Glucose-6-phosphate dehydrogenase was assayed in 10mM MgCl_2_ and 50 mM Tris maleate, pH 7.5 with 2 mM NADP+ and 4 mM glucose-6-phosphate (Sigma-Aldrich) as substrate (Shaeffer and Stanier, 1978). 6-phosphogluconate dehydrogenase was assayed in 10mM MgCl_2_ and 50 mM Tris maleate, pH 7.5 with 2 mM NADP+ and 2 mM 6-phosphogluconate (Sigma-Aldrich) as substrate (Shaeffer and Stanier, 1978). Phosphoglucomutase was assayed in 50 mM Tris-HCl, pH 8.0 and 10 mM DTT with 2 mM MgCl_2_, 10 μM glucose-1,6-bisphosphate, 1 mM NADP+, 0.2 U ml^-1^ glucose-6-phosphate dehydrogenase and 2 mM glucose-1-phosphate (Sigma-Aldrich) as substrate (Liu et al., 2013). Glycogen synthase was assayed in a three-step reaction following Suzuki et al., 2010. First ADP was generated from ADP-glucose in a reaction consisting of 50 mM Tris-HCl, pH 8.0, 20 mM DTT, 2 mM ADP-glucose (Sigma-Aldrich), 2 mg ml^-1^ oyster glycogen (Sigma-Aldrich) as substrate and the cell extract. The reaction was incubated for 20 min at 30 °C and then stopped by heat treatment at 100 °C for 2 min. Then, ATP was generated from ADP by mixing the solution 3:1 with a buffer containing 50 mM HEPES-NaOH, pH 7.5, 10 mM phosphocreatine (Sigma-Aldrich), 200 mM KCl, 10 mM MgCl_2_, and 0.5 mg ml^-1^ creatine phosphokinase (Sigma-Aldrich), and incubated for 30 min at 30 °C. The reaction was stopped by heat treatment in a 100 °C metal block for 2 min. Finally, the amount of ATP was measured by mixing the solution 3:2 with a buffer comprising of 125 mM HEPES-NaOH, pH 7.5, 10 mM glucose, 20 mM MgCl_2_, and 1 mM NADP^+^ and 5 U ml^-1^ each of hexokinase (Sigma-Aldrich) and glucose-6-phosphate dehydrogenase. All above assays were measured spectrophotometrically by monitoring the increase in absorbance at 340 nm over time using Spectramax i3 plate reader. ADP-glucose pyrophosphorylase was assayed in 50 mM HEPES, pH 8.0, 10 mM MgCl_2_ with 2 mM ATP, 2 mM 3-phosphoglycerate (Santa Cruz Biotechnology), 1 U ml^-1^ yeast inorganic pyrophosphatase (Sigma-Aldrich) and 1mM glucose-1-phosphate (Sigma-Aldrich) as substrate (Diaz-Troya et al., 2014). Production of phosphate was monitored using EnzCheck Assay (Molecular Probes) by reading absorbance at 360 nm over time in Spectramax i3 plate reader. To determine background control assays for each enzymatic activity were performed without addition of the relevant substrate.

### Glycogen measurements

Glycogen measurements were performed following Pattanayak et al., 2015 with modifications. Briefly, for each time point 10 ml of cultures were collected by filtering and freezing in liquid nitrogen. Cells were resuspended from the filter in 1 ml ice-cold methanol. Filters were discarded and cells were centrifuged at 14000 rpm for 7 minutes. The chlorophyll content was determined spectrophotometrically at 665 nm using the equation C_Chl_= A_665_ x 13.9 μg ml^-1^ (de Marsac and Houmard, 1988). Glycogen content was normalized by chlorophyll content in each sample. The pellets were resuspended in 300 μl of 40% KOH and incubated for 90 minutes at 95 °C. 200 proof ethanol was added to each extract and extracts were incubated at −20 °C overnight. Samples were centrifuged at 14000 rpm for 1h at 4 °C, the supernatants were discarded and the pellets were washed twice with ice-cold ethanol. The pellets were resuspended in 100 μl of 2 N HCL and placed in a 95 °C heat block for 30 min to break glycogen down. Samples were neutralized with 100 μl of 2 N NaOH and 50 μl of 1 M phosphate buffer, pH 7. Glycogen content was then assayed enzymatically using glucose hexokinase assay (Sigma-Aldrich) in 96-well plate at 340 nm in the Spectramax i3 plate reader. Glycogen from bovine liver (Sigma-Aldrich) was used to generate a standard curve.

## Acknowledgements

We thank Andrew Murray, Joseph Markson, Brian Zid, Alicia Darnell, Lauren Surface, Xiao-yu Zheng and James Martenson for comments on the manuscript. We thank members of the O’Shea and Denic labs for helpful discussions and constructive criticism. We thank Joseph Markson and Susan Golden for plasmids. This work was funded by the Howard Hughes Medical Institute.

---

**Figure 1 - figure supplement 1.**

**Characterization of growth and viability of the *rpaA* mutants.**

For A, C and D ten-fold serial dilutions of each culture were spotted on BG-11 medium plates and incubated for 7 days in light or for 14 days in oscillating light/dark conditions. The data are representative of three independent experiments. (A) Complementation of the *rpaA*^-^ strain with *rpaA* expressed from the neutral site NS1 rescues the viability defect in light/dark conditions. (B) Viability of the indicated strains after prolonged incubation in darkness. The survival of cells was assessed by a colony forming unit assay. Points represent the mean of two experiments with error bars displaying the standard error of the mean. (C) The *rpaA*^-^ “clock rescue” strain phenocopies the *rpaA*^-^ strain. (D) Comparison of growth of the “clock rescue” *_c_rpaA* strain and the “clock rescue” *_c_rpaA D53A* mutant, where *_c_rpaA* indicates constitutive expression of *rpaA*. In the wild type strain RpaA activates its own expression. In the *_c_rpaA* strains the *rpaA* promoter was changed to a P*0050* promoter allowing for constitutive expression of *rpaA D53A*.

**Figure 1 - figure supplement 2.**

**The *rpaA^-^* strain displays a defect in levels of adenine nucleotides in light/dark cycles.**

Quantification of relative levels of all adenine nucleotides in wild type and the *rpaA*^-^ strains grown in light/dark cycles. Points represent the mean of four experiments with error bars displaying the standard error of the mean.

**Figure 2 - figure supplement 1.**

**Expression of genes encoding enzymes involved in alternative sugar catabolism pathways is defective in the *rpaA^-^* strain in light/dark cycles.**

(A) Comparison of the normalized gene expression in the “clock rescue” and the *rpaA*^-^ “clock rescue” strains 1h after exposure to darkness. Circadian genes whose expression peaks at dawn in the wild type strain are colored yellow, dusk-peaking genes are blue and non-circadian genes are black. Fold changes are displayed graphically as diagonal lines on the plot. (B) Analysis of the functional annotations of the protein coding genes whose expression is defective in the *rpaA*^-^ “clock rescue” strain. We identified 88 protein coding genes whose expression is at least 3-fold lower in the *rpaA*^-^ “clock rescue” strain than in the control strain, as described in the Methods section. The detailed list of genes, whose expression is abrogated in the *rpaA*^-^ “clock rescue” strain, is presented in Supplementary File 1. (C) A representative photograph of a plate viability assay performed using mutants characterized by Doolittle and Singer, 1974 and Scanlan et al., 1995. The deletion mutants of indicated genes involved in alternative sugar metabolism pathways were plated as a dilution series on BG-11 agar and grown in continuous light for 7 days or under light/dark conditions for 14 days. The experiment was performed three independent times.

**Figure 3 - figure supplement 1.**

**Comparison of growth and viability of the *rpaA^-^* strain and glycogen synthesis mutants.**

(A) Comparison of growth curves of wild type, the *rpaA*^-^ strain and the deletion mutants deficient in glycogen synthesis (*glgA^-^* and *glgC^-^*) in constant light (top) and in light dark cycles (below). Points represent the mean of two experiments with error bars displaying the standard error of the mean. (B) Viability of wild type, the *rpaA*^-^ strain and strains defective in glycogen synthesis after incubation in prolonged darkness. The survival of cells was assessed by a colony forming unit assay. Points represent the mean of two experiments with error bars displaying the standard error of the mean.

**Figure 4 - figure supplement 1.**

**Characterization of strains used for the reconstitution experiments.**

Enzyme activities of glycogen phosphorylase, glucose 6-phophate dehydrogenase and 6-phosphogluconate dehydrogenase in wild type, the *rpaA^-^* and the *rpaA^-^ +enzymes* strains. Strains were incubated in BG-11 with 100 mM IPTG for 12 h in light and harvested for the assay 3 h after exposure to darkness. Error bars display the standard error of the mean of two experiments.

**Supplementary File 1.**

**Genes whose expression is abrogated in the *rpaA^-^* “clock rescue” strain.**

**Supplementary File 2.**

**Strain list.**

Cmr, chloramphenicol resistance; Gmr, gentamycin resistance; Kmr, kanamycin resistance; Ntr, nourseothricin resistance; Sp/St, spectinomycin/streptomycin resistance; NS 1, neutral site 1 (GenBank U30252)

## References

Atkinson DE. (1968). Energy charge of the adenylate pool as a regulatory parameter. Interaction with feedback modifiers. Biochemistry 7:4030–34. doi: 10.1021/bi00851a033.

Cervený J, Nedbal L. 2009. Metabolic rhythms of the cyanobacterium *Cyanothece sp*. ATCC 51142 correlate with modeled dynamics of circadian clock. J. Biol. Rhythms 24:295–303. doi: 10.1177/0748730409338367.

Chapman AG, Fall L, Atkinson DE. 1971. Adenylate energy charge in *Escherichia coli* during growth and starvation. J. Bacteriol. 108:1072–1086.

Clerico EM, Ditty, JL, Golden SS. 2007. Specialized techniques for site-directed mutagenesis in cyanobacteria. Methods Mol. Biol. 362:155–171. doi: 10.1007/978-1-59745-257-1_11.

de Marsac NT, Houmard J. 1988. Complementary chromatic adaptation: physiological conditions and action spectra. Methods Enzymol. 167:318–328.

Diamond S, Jun D, Rubin BE, Golden SS. 2015. The circadian oscillator in *Synechococcus elongatus* controls metabolite partitioning during diurnal growth. Proc. Natl. Acad. Sci. USA 112:E1916–E1925. doi: 10.1073/pnas.1504576112.

Díaz-Troya S, López-Maury L, Sánchez-Riego AM, Roldán M, Florencio FJ. 2014. Redox regulation of glycogen biosynthesis in the cyanobacterium *Synechocystis sp*. PCC 6803: analysis of the AGP and glycogen synthases. Mol. Plant. 7: 87–100. doi: 10.1093/mp/sst137.

Dodd AN, Belbin FE, Frank A, Webb AAR. 2015. Interactions between circadian clocks and photosynthesis for the temporal and spatial coordination of metabolism. Front Plant Sci. 6:245. doi: 10.3389/fpls.2015.00245.

Doolittle WF. 1979. The cyanobacterial genome, its expression, and the control of that expression. Adv Microb Physiol. 20:1–102.

Doolittle WF, Singer RA. 1974. Mutational analysis of dark endogenous metabolism in the blue-green bacterium *Anacystis nidulans*. J Bacteriol. 119:677–683.

Fu J, Xu X. 2006. The functional divergence of two *glgP* homologues in *Synechocystis sp*. PCC 6803. FEMS microbiol. lett. 260:201–209. doi: 10.1111/j.1574-6968.2006.00312.x.

Gibson DG, Young L, Chuang RY, Venter JC, Hutchison CA 3rd, Smith HO. 2009. Enzymatic assembly of DNA molecules up to several hundred kilobases. Nat Methods. 6:343–5. doi: 10.1038/nmeth.1318.

Graf A, Schlereth A, Stitt M, Smith AM. 2010. Circadian control of carbohydrate availability for growth in *Arabidopsis* plants at night. Proc. Natl. Acad. Sci. USA, 107:9458–9463. doi: 10.1073/pnas.0914299107

Green CB, Takahashi JS, Bass J. 2008. The meter of metabolism. Cell 134:728–742. doi: 10.1016/j.cell.2008.08.022

Grundel M, Scheunemann R, Lockau W, Zilliges Y. 2012. Impaired glycogen synthesis causes metabolic overflow reactions and affects stress responses in the cyanobacterium *Synechocystis sp*. PCC 6803. Microbiology 158:3032–3043. doi: 10.1099/mic.0.062950-0.

Knoop H, Zilliges Y, Lockau W, Steuer R. 2010. The Metabolic Network of *Synechocystis sp*. PCC 6803: Systemic Properties of Autotrophic Growth. Plant Physiology 154, 410–422. doi: 10.1104/pp.110.157198.

Lambert G, Chew J, Rust MJ. 2016. Costs of Clock-Environment Misalignment in Individual Cyanobacterial Cells. Biophysical Journal 111:883–891. doi: 10.1016/j.bpj.2016.07.008.

Liu L, Hu HH, Gao H, Xu XD. 2013. Role of two phosphohexomutase genes in glycogen synthesis in *Synechocystis sp*. PCC6803. Chinese Science Bulletin 58(36): 4616–4621. doi: 10.1007/s11434-013-5958-0

Markson JS, Piechura JR, Puszynska AM, O’Shea EK. 2013. Circadian control of global gene expression by the cyanobacterial master regulator RpaA. Cell 155: 1396–1408. doi: 10.1016/j.cell.2013.11.005.

Nishiwaki T, Satomi Y, Kitayama Y, Terauchi K, Kiyohara R, Takao T, Kondo T. 2007. A sequential program of dual phosphorylation of KaiC as a basis for circadian rhythm in cyanobacteria. EMBO J. 26: 4029–37. doi: 10.1038/sj.emboj.7601832.

Ouyang Y, Andersson CR, Kondo T, Golden SS, Johnson CH. 1998. Resonating circadian clocks enhance fitness in cyanobacteria. Proc. Natl. Acad. Sci. USA 95:8660–8664.

Pattanayak GK, Phong C, Rust MJ. 2014. Rhythms in energy storage control the ability of the cyanobacterial circadian clock to reset. Curr. Biol. 24(16):1934–8. doi: 10.1016/j.cub.2014.07.022.

Pattanayak GK, Lambert G, Bernat K, Rust MJ. 2015. Controlling the Cyanobacterial Clock by Synthetically Rewiring Metabolism. Cell Reports 13(11):2362–2367. doi: 10.1016/j.celrep.2015.11.031

Rugen M, Bockmayr A, Steuer R. 2015. Elucidating temporal resource allocation and diurnal dynamics in phototrophic metabolism using conditional FBA. Sci. Rep. 5:15247. doi:10.1038/srep15247

Rust MJ, Golden SS, O’Shea EK. 2011. Light-driven changes in energy metabolism directly entrain the cyanobacterial circadian clock. Science 331:214–217. doi: 10.1126/science.1197243.

Rust MJ, Markson JS, Lane WS, Fisher DS, O’Shea EK. 2007. Ordered phosphorylation governs oscillation of a three-protein circadian clock. Science 318:809–812. doi: 10.1126/science.1148596.

Scanlan DJ, Sundaram S, Newman J, Mann NH, Carr NG. 1995. Characterization of a *zwf* mutant of Synechococcus sp. strain PCC 7942. J Bacteriol. 177:2550–53.

Schaeffer F, Stanier RY. 1978. Glucose-6-phosphate dehydrogenase of *Anabaena* sp. Kinetic and molecular properties. Arch. Microbiol. 116:9–19. doi: 10.1007/BF00408728.

Smith AJ. 1983. Modes of cyanobacterial carbon metabolism. Ann. Microbiol. 134B: 93–113. doi: 10.1016/S0769-2609(83)80099-4.

Suzuki E, Ohkawa H, Moriya K, Matsubara T, Nagaike Y, Iwasaki I, Fujiwara S, Tsuzuki M, Nakamura Y. 2010. Carbohydrate metabolism in mutants of the cyanobacterium *Synechococcus elongatus* PCC7942 defective in glycogen synthesis. Appl. Environ. Microbiol. 76:3153–3159. doi: 10.1128/AEM.00397-08

Takai N, Nakajima M, Oyama T, Kito R, Sugita C, Sugita M, Kondo T, Iwasaki, H. 2006. A KaiC associating SasA-RpaA two-component regulatory system as a major circadian timing mediator in cyanobacteria. Proc. Natl. Acad. Sci. USA 103: 12109–14. doi: 10.1073/pnas.0602955103.

Taniguchi Y, Takai N, Katayama M, Kondo T, Oyama T. 2010. Three major output pathways from the KaiABC based oscillator cooperate to generate robust circadian *kaiBC* expression in cyanobacteria. Proc. Natl. Acad. Sci. USA 107:3263–68. doi: 10.1073/pnas.0909924107.

Taton A, Unglaub F, Wright NE, Zeng WY, Paz-Yepes J, Brahamsha B, Palenik B, Peterson TC, Haerizadeh F, Golden SS, Golden JW. 2014. Broad-host-range vector system for synthetic biology and biotechnology in cyanobacteria. Nucleic Acids Res. 42:e136. doi: 10.1093/nar/gku673.

Teng SW, Mukherji S, Moffitt JR, de Buyl S, O’Shea and EK. 2013. Robust circadian oscillations in growing cyanobacteria require transcriptional feedback, Science 340:737–740. doi: 10.1126/science.1230996.

Tu BP and McKnight SL. 2006. Metabolic cycles as an underlying basis of biological oscillations. Nat. Rev. Mol. Cell Biol. 7:696–701. doi: 10.1038/nrm1980.

Vijayan V, Zuzow R, O’Shea EK. 2009. Oscillations in supercoiling drive circadian gene expression in cyanobacteria. Proc. Natl. Acad. Sci. USA 106:22564–68. doi: 10.1073/pnas.0912673106.

Woelfle MA, Ouyang Y, Phanvijhitsiri K, Johnson CH. 2004. The adaptive value of circadian clocks: an experimental assessment in cyanobacteria. Curr. Biol. 14:1481–1486. doi: 10

